# Quantitative PCR provides a simple and accessible method for quantitative microbiome profiling

**DOI:** 10.1101/478685

**Authors:** Ching Jian, Panu Luukkonen, Hannele Yki-Järvinen, Anne Salonen, Katri Korpela

## Abstract

The use of relative next generation sequencing (NGS) abundance data can lead to misinterpretations of microbial community structures as the increase of one taxon leads to concurrent decrease of the other(s). To overcome compositionality, we provide a quantitative NGS solution, which is achieved by adjusting the relative 16S rRNA gene amplicon NGS data with quantitative PCR (qPCR-based) total bacterial counts. By comparing the enumeration of dominant bacterial groups on different taxonomic levels in human fecal samples using taxon-specific 16S rRNA gene-targeted qPCR we show that quantitative NGS is able to estimate absolute bacterial abundances accurately. We also observed a higher degree of correspondence in the estimated microbe-metabolite relationship when quantitative NGS was applied. Being conceptually and methodologically analogous to amplicon-based NGS, our qPCR-based method can be readily incorporated into the standard, high-throughput NGS sample processing pipeline for more accurate description of interactions within and between the microbes and host.

## Main text

Microbiota datasets arising from NGS are compositional in nature: the relative abundances of the taxa always sum up to 1 (1). Since the changes of components are mutually dependent, erroneous conclusions may occur if data are analyzed using traditional statistical methods. Compositionality particularly hampers the interpretation of microbiota changes in longitudinal studies, such as interventions. If a single taxon increases or decreases in relative abundance, the relative abundances of other taxa will show a corresponding, opposite change. It becomes impossible to determine which taxon was truly affected by an intervention (Supplementary Fig. S1). Contrary to the speculation that compositionality is dismissible in high complexity environments (2), our simulations revealed that the compositionality effects may lead to extensive false positive findings in samples containing complex microbial communities (e.g., gut) as well as samples with low diversity (e.g., vaginal microbiome) (Supplementary Fig. S2 and S3). Complex analytical methods have been developed in an effort to mitigate the effect of mutual dependence of component changes in compositional microbiota sequencing datasets (2, 3). However, in addition to the technical problems of compositional data analysis, there is reason to believe that the absolute abundances of bacteria are a biologically meaningful measure and relying solely on relative abundances results in the omission of important information on the interactions of different taxa with each other and the host (4). Absolute quantification of microbial abundances based on flow cytometry has been applied to complement amplicon sequencing and shown to more precisely describe the temporal dynamics of the microbial community in an engineered freshwater ecosystem (5). Vandeputte et al. (6) developed an integrative workflow that combines flow cytometry-based bacterial enumeration and NGS for fecal samples. Also spike-in bacteria (7) or DNA (8) has been used for the purpose of quantification of microbiota NGS profiles. For fungi, also qPCR has been employed to transform the relative abundance data from pyrosequencing to absolute values (9). Notwithstanding the increased awareness of compositionality and the recent advances in quantitative microbiome profiling, the majority of human microbiome studies to date have not adopted any countermeasures. We believe the practicability and accessibility of many previously proposed solutions represent obstacles for the wider adaptation of quantitative microbiome profiling in human microbiome research. In particular, the flow cytometry-based approach requires considerable expertise for reproducible results, considering flow cytometric enumeration of microbial cells was initially restricted to pure cultures (10) and still remains challenging when performed in complex matrices (11).

To render quantitative microbiome profiling from NGS data easy and accessible, we devised and validated a 16S rRNA gene-targeted qPCR-based quantification method. qPCR has been widely used for accurate and targeted quantification of specific bacterial groups and species especially until the era of NGS. Here, we first quantified total bacteria using universal bacterial primers (12) by qPCR (Supplementary Methods) in 114 adult fecal DNA samples that have been analyzed for microbiota composition using Illumina MiSeq for 16S rRNA gene amplicon sequencing (13). The qPCR threshold cycle (Ct) values were converted to the estimates of bacterial genomes present in 1 g of feces (total bacterial counts), which were used to estimate absolute abundance of the NGS-detected taxa (Fig. 1) by a simple equation:

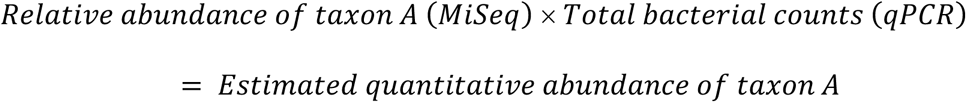

**Figure 1:**
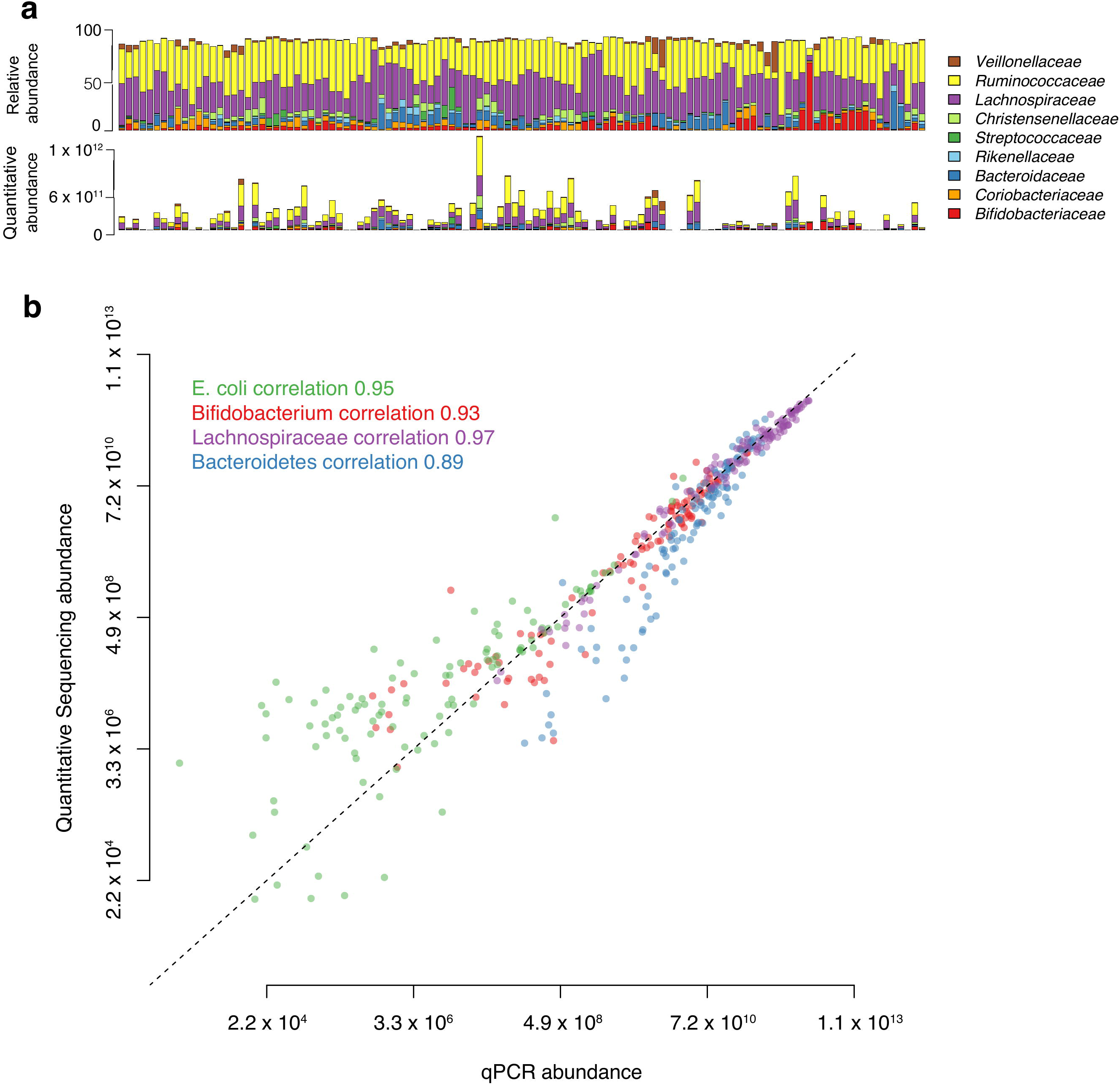
Relative microbiota profiles translated into quantitative microbiota profiles using quantitative NGS. **a**, Comparison of relative abundance and quantitative sequencing abundance of dominant bacterial families in 114 fecal samples. **b**, Correlation between the qPCR abundance (16S rRNA gene copies per g feces) and the quantitative sequencing abundance of four taxa representing the species, genus, family and phylum levels. The dashed line shows the expected 1:1 correspondence.

Next, we validated the estimated quantitative abundances of four representative taxa by qPCR using the taxon-specific primers (Supplementary Table 1) for the phylum Bacteroidetes, *Clostridium* cluster XIVa (family *Lachnospiraceae*), genus *Bifidobacterium* and *Escherichia coli* species. Absolute quantification was achieved by using standard curves. We found near-perfect correlations between the estimated quantitative abundances and qPCR abundances in all tested taxa (Fig. 1b), indicating that qPCR-adjusted NGS (hereafter termed quantitative NGS) profiles are able to reflect absolute bacterial abundances in feces as accurately as taxon-specific qPCR. The correspondence decreases at the very low end of the abundance range, likely due to the relatively lower PCR amplification efficiency and increased stochasticity of low abundance taxa (14). The applied library preparation method (dual index TruSeq-tailed 1-step amplification (15)) causes a slight underestimation of Bacteroidetes abundance (unpublished data), explaining the underestimation observed for this phylum compared to qPCR (Fig. 1b).

To determine whether the quantitative NGS data can be used to more precisely estimate the abundance of specific gut microbiota functions, we correlated quantitative NGS abundance of butyrate-producing bacteria to the abundance of the butyryl-CoA:acetate CoA-transferase gene determined by qPCR (16). As expected, quantitative NGS data explained a much larger proportion of the variation in butyryl-CoA gene abundance than the relative abundance of the butyrate producers (variance explained 0.47 versus 0.23, Fig. 2).

**Figure 2:**
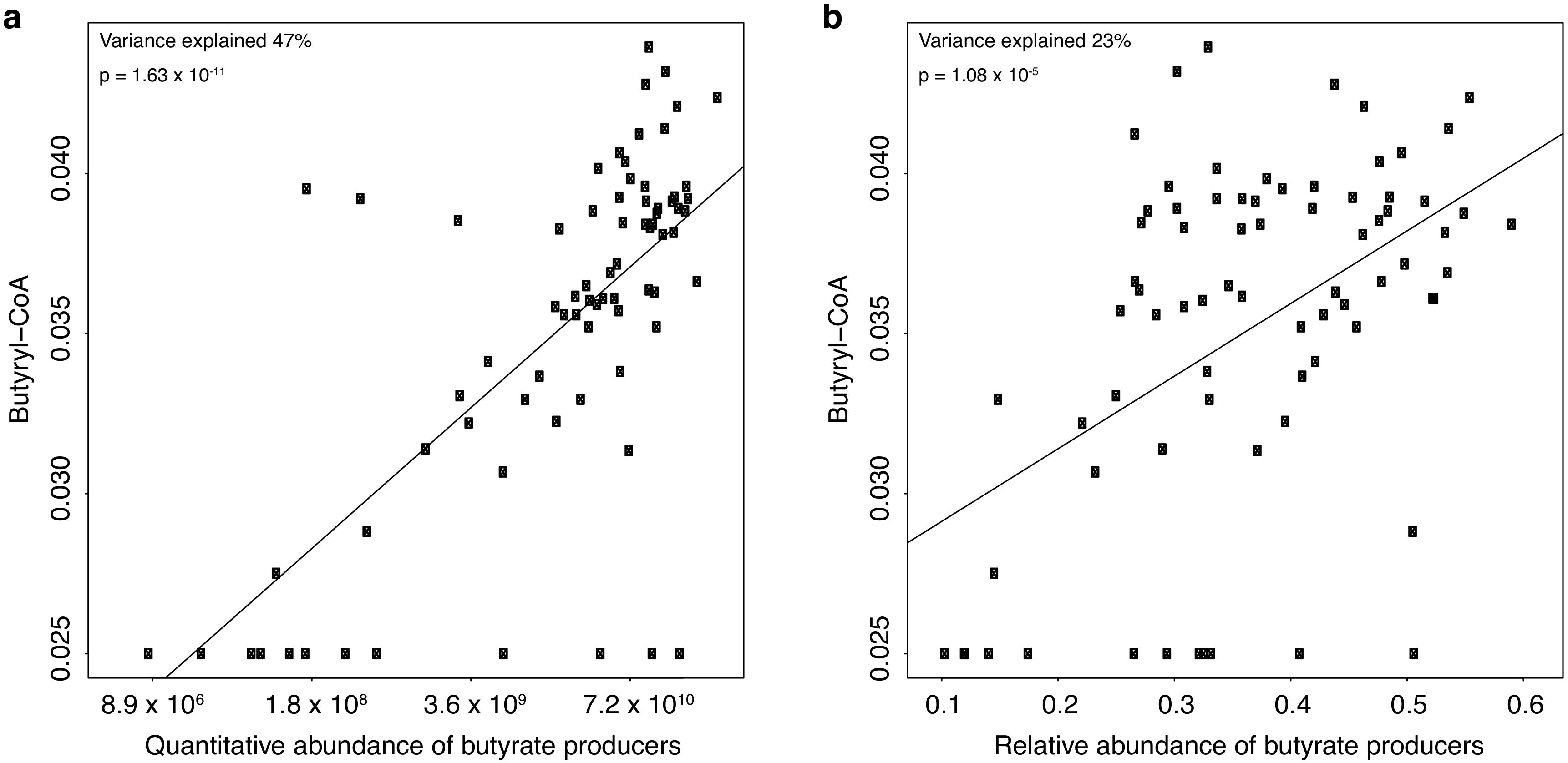
Association between the abundance of the butyryl-CoA:acetate CoA-transferase gene, measured by qPCR, and the abundance of butyrate producers. **a**, Sequencing-based quantitative abundance. **b**, Relative abundance. The variance explained and the p-value are from a linear regression.

We contend that our qPCR-based quantitative NGS enjoys conceptual and practical benefits over other alternatives for quantification of the bacterial load in the NGS samples. First and foremost, the same DNA extract serves as the starting material both for qPCR and NGS, making qPCR easy to implement in the workflow for high-throughput analysis of up to thousands of microbiome samples. Second, unlike flow cytometry that counts cells, qPCR and NGS both target bacterial DNA, including extracellular DNA derived from lysed bacteria. Extracellular DNA can be intrinsic or result from the differential lysis of Gram-positive and negative bacteria during the common freeze-thawing prior to fecal DNA extraction. As the 16S profiles from the gut appear very different for intracellular and extracellular DNA (17), qPCR is expected to reflect the NGS-targeted community structure both quantitatively and qualitatively more closely than flow cytometry. Third, qPCR is cost-effective and more accessible as the laboratory settings, machinery and reagents are essentially the same as those needed for preparing the NGS libraries. While PCR-based methods can be potentially biased e.g. due to inadequate DNA extraction or primer coverage, these factors play a similar role in the NGS itself (18). Simplicity is another strength, as no spikes or other exogenous controls are needed. Lastly, our quantitative microbiome profiling approach is compatible with any downstream bioinformatic pipelines, since it requires no complicated transformation or computation.

In conclusion, we caution against the analysis of microbiome NGS data solely relying on relative abundances, since compositionality may skew biological inferences from microbiota studies *per* our simulation data as well as the previously published studies. Although relative taxon abundance can be indicative at times, absolute quantification is necessary for obtaining a comprehensive understanding of the dynamics and interactions of the microbiome. To facilitate the standard use of quantitative microbiome data, we propose a qPCR-based quantitative NGS approach that entails a single additional amplification reaction and can be integrated into an NGS workflow seamlessly. While our proposed approach does not solve the limitations and biases inherent to the NGS technology, we believe our reductionist solution to compositionality offers researchers improved biological insight from their microbiota sequencing datasets.

## Supporting information

## Acknowledgements

We thank Professor Willem de Vos for comments on the manuscript.

## Contributions

C.J. performed the laboratory analyses and drafted the manuscript.

A.S. supervised the laboratory work, reviewed and edited the manuscript.

H.Y.-J. and P.L. designed and conducted the clinical trial from where the fecal samples originated.

K.K. conceived the study, performed the statistical analyses, reviewed and edited the manuscript.

