## Supplementary material for "Quantitative PCR provides a simple and accessible method for quantitative microbiome profiling"

This document contains the following information:

Supplementary Methods

Supplementary Figures 1-3

Supplementary Table 1

Supplementary References

**Supplementary Methods**

*Study subjects and fecal sample collection*

The study cohort consisting of 38 adult human subjects has been described previously (1). The study protocol has been approved by the Medical Ethical Committees of the Hospital District of Helsinki and Uusimaa and HUCH. For the current study, we additionally included follow-up samples collected after the trial, amounting to a total of 114 samples. Fecal samples were self-collected and stored at -20°C, and then transferred to the long-term storage at -80°C within 1 day.

*Bacterial DNA extraction*

Bacterial DNA was extracted from fecal samples using a modified version of repeated bead beating (2) that efficiently extracts bacterial DNA from both Gram-positive and negative cocci (3). Briefly, immediately after thawing, 0.125 g of feces were weighted and added into 2.0 ml screw-up tubes pre-filled with 0.25 g of 0.1 mm zirconia beads and 3 of 3 mm glass beads. Fecal samples were re-suspended to 0.5 ml of lysis buffer (500 mM NaCl, 50 mM Tris-HCL (pH 8), 50 mM EDTA, 4 % SDS). Two successive rounds of bead beating were done using a FastPrep®-24 instrument (MP Biomedicals, Santa Ana, CA, USA) at 5.5 $m/s^{-1}$. The lysate fraction produced from the first round of bead beating was collected before the second round to minimize DNA shearing (2). Each round of bead beating was followed by a 15-min incubation period at 95°C to further enhance the lysis. After precipitation of DNA, the DNA was further purified by using the QIAamp DNA Mini Kit columns (Qiagen, Hilden, Germany). The purified DNA was quantified for DAN concentration using a Qubit® fluorometer (Invitrogen, CA, USA) before storing at -20° C until further use. All the fecal samples were processed within 10 days.

*16S rRNA gene sequencing*

Illumina MiSeq paired-end sequencing of hypervariable V3-V4 regions of the 16S rRNA gene was performed according to the manual from Illumina with a slight modification where dual index TrusSeq-tailed 1-step amplification (4) was used for library preparation. The detailed protocol for library preparation has been described (1). The pooled libraries were sequenced with an Illumina MiSeq instrument using paired end 2 × 300 b.p. reads and a MiSeq v3 reagent kit with 5% PhiX as spike-in. The sequencing was carried out at the sequencing unit of the Institute for Molecular Medicine Finland (FIMM), Helsinki, Finland.

*Sequencing data processing and analysis*

The preprocessing was done in the R package *mare* (5). Only the forward reads were used in order to minimize phylogenetically biased data as recommended (5). The forward reads were truncated to length of 150 bases with *mare*'s "ProcessReads" command. We used default settings for minimum quality score and maximum expected errors. Reads with prevalence below 0.01% were removed. Chimera removal and dereplication of the reads was done using USEARCH8 (6). Truncated, filtered and dereplicated reads were annotated using the Silva database (7).

*Quantitative PCR*

Quantification of total bacteria, specific taxa and butyrate production capacity was carried out by qPCR using a BioRad iCycler iQ thermal cycler system (BioRad, Hercules, CA) with HOT FIREPol® EvaGreen® qPCR Mix Plus (Solis BioDyne, Tartu, Estonia). A list of primers and references used in the present study is summarized in Supplementary Table 1.

For bacterial enumeration, total bacteria, Clostridium cluster XIVa and Bacteroidetes were quantified using 0.5 ng of fecal DNA, for the less abundant *Bifidobacterium* and *E. coli* groups 25 ng DNA/reaction was used. Detailed information on the PCR conditions has been described previously (2, 8). The 10-log-fold standard curves ranging from $10^2$ to $10^7$ copies were produced using the full-length amplicons of 16S rRNA gene of appropriate reference organisms (8) (*Ruminococcus productus, Bacteroides fragilis, Bifidobacterium longum and Escherichia coli*

*DSM 6897)* to convert the threshold cycle (Ct) values into the average estimates of target bacterial genomes present in 1 g of feces (copy numbers/g of wet feces) in each assay.

For quantification of butyrate production capacity of the microbiota, the butyryl-CoA:acetate CoA-transferase gene was quantified by qPCR as described (9), and the output values were converted based on comparative Ct method (10). The results were correlated to NGS-based abundance of the dominant butyrate-producing genera *Subdoligranulum, Faecalibacterium, Anaerostipes, Butyrivibrio, Coprococcus, Blautia* and *Roseburia/Eubacterium rectale* (9).

All qPCR assays were performed in triplicate. Precautions were taken to ensure that the data from each triplicate fall within 0.5 threshold cycle (Ct), and clear outliers (>2 standard deviations) were removed before calculating average Ct of each sample. Melting curves and non-template controls were used to assess run reliability. There was no detectable amplification arising from non-template controls in any of the assays. The amplification efficiencies of all qPCR assays ranged from 91% to 98%.

*Data availability*

All data are available from the corresponding author upon request.

**Supplementary Table**

Supplementary table 1: List of primers used in this study.

| Target | Primer sequence | Original reference |
|---|---|---|
| Bacteria (total bacteria) | F: 5'-TCCTACGGGAGGCAGCAGT-3'<br>R: 5'-GGACTACCAGGGTATCTAATCCTGTT-3' | (11) |
| *Bacteroides-Prevotella-Porphyromonas* | F: 5'-GGTGTCGGCTTAAGTGCCAT-3'<br>R: 5'-CGGA(C/T)GTAAGGGCCGTGC-3' | (8) |
| *Bifidobacterium* spp. | F: 5'-TCGCGTC(C/T)GGTGTGAAAG-3'<br>R: 5'-CCACATCCAGC(A/G)TCCAC-3' | (8) |

| | | |
|---|---|---|
| *Clostridium coccoides-Eubacterium rectale* group (*Lachnospiraceae*) | F: 5'-CGGTACCTGACTAAGAAGC-3' | (8) |
| | R: 5'-AGTTT(C/T)ATTCTTGCGAACG-3' | |
| *Escherichia coli* subgroup (*E. coli*, *Hafnia alvei* and *Shigella* spp.) | F: 5'-GTTAATACCTTTGCTCATTGA-3' | (12) |
| | R: 5'-ACCAGGGTATCTAATCCTGTT-3' | |
| Butyryl-CoA CoA transferase | F: 5'-GCIGAICATTTCACITGGAAYWSITGGCAYATG-3' | (9) |
| | R: 5'-CCTGCCTTTGCAATRTCIACRAANGC-3' | |

### Supplementary Figures

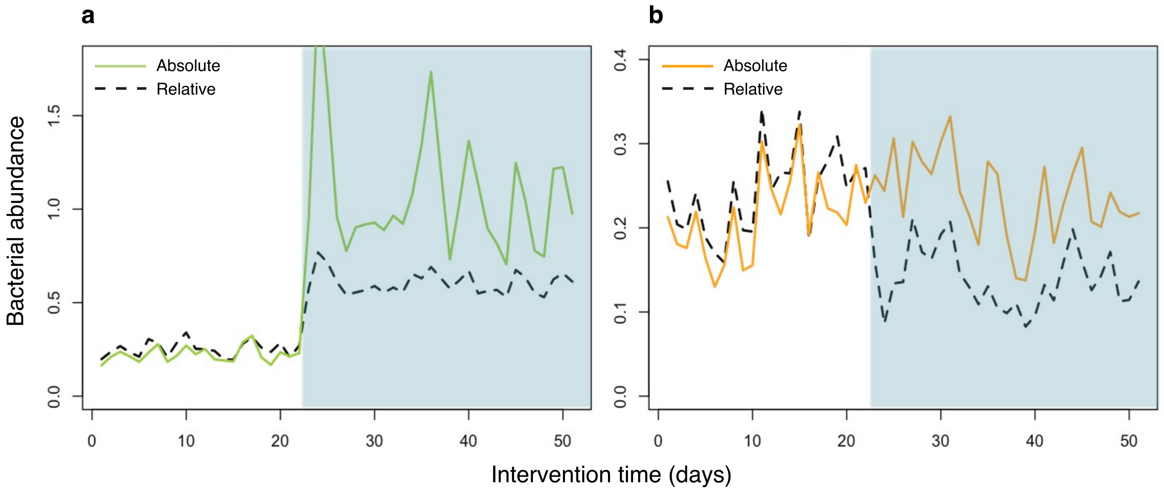


**Supplementary Figure S1: Compositionality leading to false positive discoveries.** To demonstrate the effect of compositionality on interpretation of microbiota NGS data, an intervention was simulated where one single taxon was increased in abundance. The simulation was conducted in absolute abundances, which were converted to relative abundances for data analysis. **a**, A single taxon (green solid line) was positively affected by the intervention (shaded area), which remained true when converted to relative abundance (black dashed line). **b**, Other taxa, here represented by a single taxon (orange solid line), were not affected by the intervention. However, their relative abundances (black dashed line) show a negative impact of the intervention, due to the increase in the relative abundance of affected taxon in **a**.

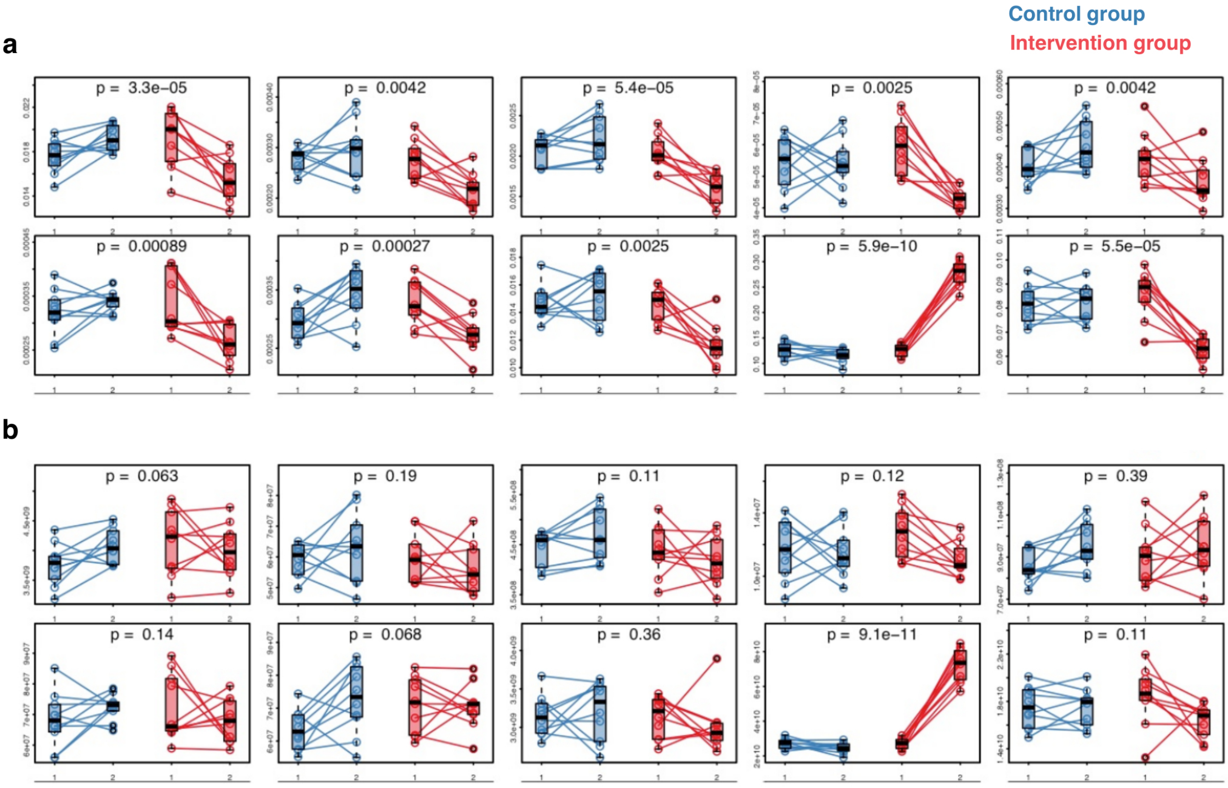


**Supplementary Figure S2: Selected results of a simulated intervention in a complex community (91 taxa).**

To demonstrate the effect of compositionality on interpretation of microbiota NGS data, another intervention was simulated where one single taxon was increased in abundance in the treatment group (n=10) and nothing changed in the control group (n=10). The panels show the 10 most significantly affected taxa (significantly different change between the control and the treatment group) based on relative abundance in the simulated intervention. Each box shows one taxon, in the same order for relative (**a**) and absolute (**b**) data. The change in the abundance of each taxon from baseline to the post-intervention sample was calculated for each individual and the significance of the difference between the treatment and the control groups was tested using analysis of variance (ANOVA). **a**, Relative data. While only one taxon was actually affected by the intervention, 62 out of 91 taxa (67%) were significantly different between the groups at p-value cut-off 0.05, and 19 taxa (21%) at p-value cut-off 0.005. **b,** Absolute data. While one taxon was affected by the intervention, additionally only 3 false positives (3%; not among the shown top 10 taxa based on the relative data) were identified with p-value cut-off 0.05 when analyzing the absolute abundances due to random variation added in the simulation, and no false positives were detected at the 0.005 level.

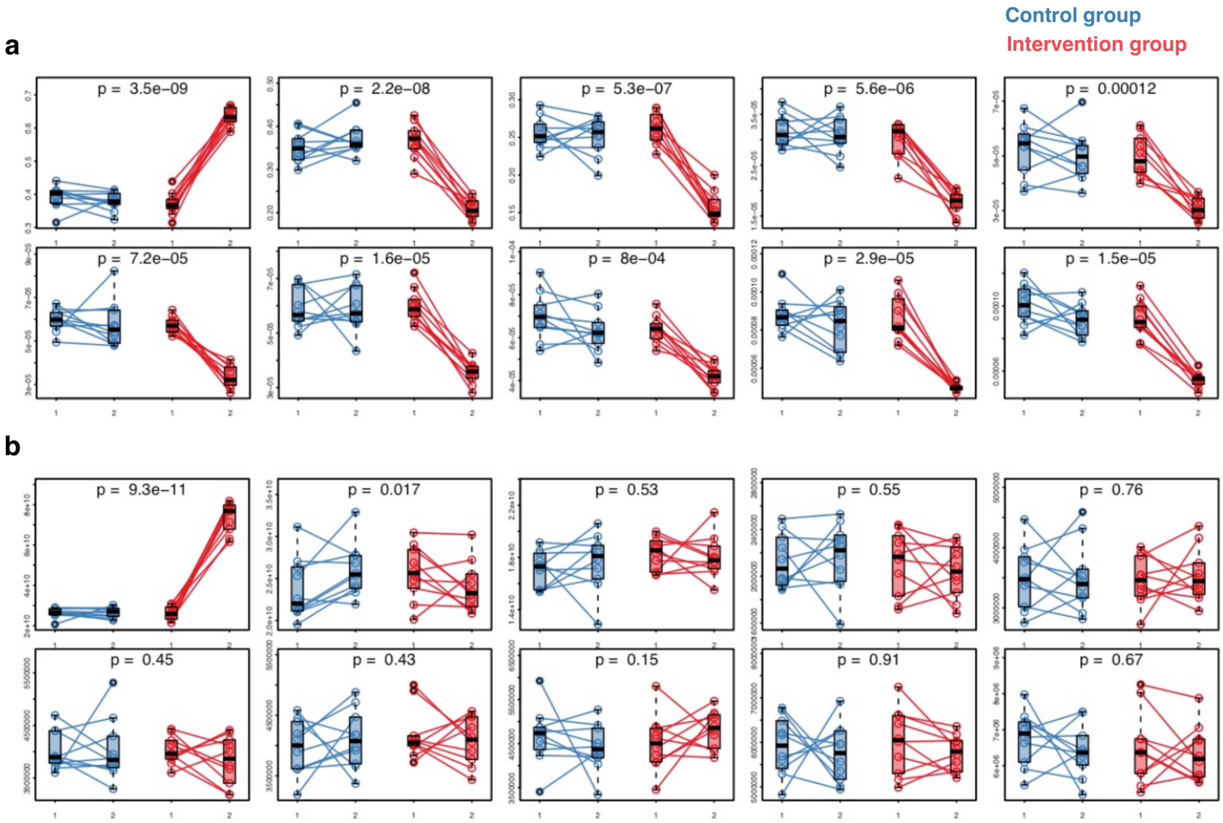


**Supplementary Figure S3: Results of a simulated intervention in a simple community (10 taxa).**

In the same simulated intervention, the compositionality problem was even more debilitating when the community is simple. **a**, When the relative abundance data were analyzed, all taxa appeared significantly affected by the intervention at strict p-value cut-off 0.005. **b**, Absolute abundance data show correctly that only one taxon was actually affected by the intervention (p-value < 0.005). No false positives were detected.
